# Restoring Synaptic Balance in Schizophrenia: Insights from a thalamo-cortical conductance-based model

**DOI:** 10.1101/2024.11.13.623344

**Authors:** Lioba C S Berndt, Krish D Singh, Alexander D Shaw

## Abstract

The dysconnectivity hypothesis of schizophrenia suggests that atypical, aberrant neural communication underlies the disorder’s diverse symptoms. Building on this framework, our study introduces a novel approach to understanding schizophrenia and exploring potential ways to adjust neural activity through synaptic restoration. Using a combination of magnetoencephalography data and dynamic causal modeling, we identified specific synaptic disturbances in schizophrenia patients, including increased NMDA receptor-mediated excitation in superficial pyramidal neurons and reduced GABA-B receptor-mediated inhibition between interneurons and pyramidal cells. These findings reveal a critical imbalance in excitation and inhibition within thalamo-cortical circuits, manifesting as altered gamma and alpha oscillations. The cornerstone of our research is an in silico synaptic restoration analysis, which demonstrates that targeted modifications to AMPA, NMDA, GABA-A, and GABA-B receptor-mediated connections can recalibrate altered neural activity in schizophrenia, aligning it with healthy control patterns. This restoration approach not only highlights the complex nature of synaptic dysfunction in the disorder but also identifies specific pathways as potential therapeutic targets, offering new avenues for investigating schizophrenia’s diverse symptomatology.

## 1 Introduction

Schizophrenia is a severe mental disorder characterised by a range of symptoms, including hallucinations, delusions, disorganised thinking, and impaired social functioning. Despite decades of research, the precise neurobiological mechanisms underlying schizophrenia remain elusive. To gain a deeper understanding of the pathology, there is a pressing need for techniques that allow for the *in vivo* assay of neurobiological mechanisms and cortical circuitry in patients. Traditional methods often rely on post-mortem tissue analysis [55, 33] or non-invasive imaging [50], which provide valuable insights but are limited in their ability to infer on inter-population (c.f. inter-laminar and thalamo-cortical) connectivity and receptor dynamics underwriting the schizophrenia syndrome.

One solution to this micro-macro schism is through computational models of neuronal circuitry – as dynamical systems or dynamic causal models – which offer a unique opportunity to glean insights into intricate synaptic processes and receptor dynamics by estimating these parameters from non-invasive M/EEG data [21]. Specifically, parameterised generative models based upon established biophysical neuronal equations such as the Hodgkin-Huxley model [34] can be inverted on empirical data features such as power spectral densities or event-related responses. Model inversion involves estimating the parameters of the underlying equations of motion – which correspond to the strengths of inter-population connectivities and receptor dynamics from empirical data.

Computational modelling of this sort permits inferences on brain dynamics at the same spatial scale on which we understand drug mechanisms and, crucially, the same scale at which many neuropathological hypotheses of schizophrenia exist [21]. As such, computational models may serve as tools [20] for testing hypotheses about impaired receptor systems or connectivity and help identify targets for pharmacological treatment.

### 1.1 Theoretical Framework: Schizophrenia and Dysconnectivity

Among the leading neurobiological models of schizophrenia, the *dysconnectivity hypothesis* has gained considerable attention [74]. It proposes that schizophrenia is fundamentally a disorder of altered connectivity within and between major brain networks. This hypothesis combines earlier models such as the *dopamine hypothesis* [36], which primarily focused on dopaminergic dysregulation as the root cause of schizophrenia’s positive symptoms, and the *glutamate hypothesis* [71], which implicates *NMDA receptor hypofunction* in broader cognitive and negative symptoms.

The dysconnectivity hypothesis emphasizes that it is not just isolated neurotransmitter dysfunction but rather aberrant coordination of brain regions that lies at the core of schizophrenia [74]. This disruption manifests both in cortico-cortical interactions [74] and cortico-subcortical circuits, particularly those involving the thalamus, striatum, and hippocampus [82, 37, 59]. The thalamo-cortical circuit plays a critical role in sensory processing and cognitive integration, and disruptions in this pathway have been implicated in the sensory gating deficits and cognitive disorganization that characterize schizophrenia [4, 5].

Importantly, schizophrenia is also associated with disturbances in neural oscillations, particularly in the *gamma-band* (30–80 Hz) range [64, 67, 31, 11]. Studies consistently show reduced gamma power and altered peak frequencies in individuals with schizophrenia. This oscillatory impairment aligns with the excitation-inhibition imbalance hypothesis [38], which posits that disruptions in the balance between excitatory and inhibitory neurotransmission—specifically between glutamatergic and GABAergic systems—are key to the disorder’s pathophysiology.

GABA, the primary inhibitory neurotransmitter in the brain, is central to this balance. GABAergic dysfunction contributes to impaired inhibitory control. The interplay between GABAergic inhibition and NMDA receptor-mediated excitation maintains the stability of thalamo-cortical circuits. Dysfunctions in both systems are believed to contribute to the broader network-level disruptions observed in schizophrenia, particularly in gamma-band oscillations and connectivity [15, 72].

### 1.2 Study Goals and Restoration Analysis

This study builds upon the findings of Shaw et al. (2020) [67], who investigated visual processing in schizophrenia using MEG. They demonstrated that schizophrenia patients showed reduced visual gamma frequency (which correlated with negative symptom severity) and impaired orientation discrimination performance. Using Dynamic Causal Modeling, they found reduced local synaptic connections in schizophrenia, with local inhibition correlating negatively with negative symptom severity. Crucially, they showed that the effective connectivity between inhibitory interneurons and superficial pyramidal cells predicted orientation discrimination performance, highlighting the role of inhibitory circuits in perceptual processing.

Their analysis employed a convolution-based Canonical Micro-circuit (CMC) model with four interconnected cell populations (layer 2/3 and 5/6 pyramidal cells, layer 4 spiny-stellate cells, and multi-layer inhibitory interneurons) [65]. However, without the specificity of conductance-based equations of motion, this simplified model architecture couldn’t determine which specific receptor systems drove the observed differences. This limitation is crucial because both GABAergic and NMDA receptor systems are implicated in schizophrenia pathophysiology [15, 72] and represent important potential intervention targets. Understanding these differences requires consideration of broader cortical-subcortical circuits involved in sensory processing. The thalamus plays a central role in relaying and modulating sensory information to the visual cortex [69, 40]. Dysfunction within thalamocortical circuits has been linked to abnormal oscillatory activity and impairments in sensory and cognitive functions in schizophrenia [70, 25, 17, 52].

To address these limitations, we applied a thalamocortical model, which explicitly incorporates both specific receptor systems and thalamocortical interactions, offering deeper insight into the mechanisms underlying the observed oscillatory differences in schizophrenia. We further performed synaptic restoration analysis to determine the parameter changes necessary to bring the EEG power spectrum density (PSD) of the schizophrenia group to align with the ones of healthy controls.

Our primary objectives were to:

1. **Identify the optimal model architecture** capable of explaining the observed MEG data in the frequency domain, focusing on alterations in thalamo-cortical connectivity.
2. **Quantify the inter-population connectivity** differences between the schizophrenia group and healthy controls.
3. **Examine the relationship between connectivity patterns and negative symptoms** in schizophrenia, exploring how altered neural dynamics may correlate with clinical outcomes.
4. Perform a **synaptic restoration analysis** to determine the synaptic and connectivity changes necessary to shift the models of individuals with schizophrenia toward those of healthy controls.

The restoration analysis plays a central role in this study by offering a pathway to understand how neural circuits in schizophrenia might be “corrected” to resemble the functional patterns of healthy individuals. Through this process, we aim to simulate the synaptic adjustments—across key neurotransmitter systems—that could restore balanced connectivity, providing insights into potential therapeutic targets. This approach extends beyond simply identifying dysfunction; it offers a framework for ***restoring neural synchrony*** and mitigating the cognitive and perceptual disturbances central to schizophrenia.

## 2 Methods and Materials

### 2.1 Study Design & Participants

**Table 1:** Demographic characteristics of study participants. This table presents the number of participants, mean age, and standard deviation of age for control and schizophrenia (SZ) groups, further subdivided by gender. The rightmost column shows p-values from independent t-tests comparing age between SZ and control groups overall and within each gender. No significant age differences were found between SZ and control groups.

| Group | Number | Mean Age (y) | SD Age (y) | Independent *T*-Test<br>SZ Vs Controls (*P* value) |
| --- | --- | --- | --- | --- |
| All controls | 29 | 41.1 | 10.1 | 0.16 |
| All SZ | 27 | 44.7 | 8.5 |  |
| Female controls | 11 | 40.2 | 12.1 | 0.3 |
| Female SZ | 8 | 45.9 | 8.0 |  |
| Male controls | 18 | 41.6 | 9.4 | 0.4 |
| Male SZ | 19 | 44.2 | 8.8 |  |

This study utilizes data originally collected and analyzed by Shaw et al. (2020) [67]. For context, we provide a brief overview of the original study’s methodology. The study included 28 individuals diagnosed with schizophrenia and 30 controls. Recruitment for the schizophrenia participants was conducted through the Cognition in Psychosis study by the Schizophrenia Working Group of the Psychiatric Genomics Consortium [57, 43]. Controls were sourced mainly through advertisements on the Cardiff University Noticeboard and opportunistically from CUBRIC at Cardiff University. Ethical approval for the study was granted by both the South East Wales NHS Ethics Board and the Ethics Board of the School of Psychology at Cardiff University.

The patient group inclusion criteria included a DSM-IV diagnosis of schizophrenia [10]. Participants were required to be aged between 16 and 75 years, with English as their first language, and possessing either normal or corrected-to-normal vision. The healthy control group was subjected to similar age and language requirements, and also needed to have normal or corrected vision.

All individuals were assessed to ensure their capability to provide informed consent. Screening for current drug or alcohol abuse was conducted using the MINI interview [68], with exclusion criteria including any reported harmful use. All participants were required to refrain from alcohol or drug use for 48 hours prior to testing.

Common exclusion criteria for all participants included a diagnosis of epilepsy, any major neurological incidents such as a severe head injury or stroke, and the presence of any metal in the body. Additionally, healthy controls were excluded if they or a first-degree relative had been previously diagnosed with a mental health condition, particularly affective or psychotic disorders, according to the MINI interview used for control assessments. Detailed information about inclusion procedure can be found in our previous paper [67].

### 2.2 Visual Gamma

In this study, we implemented an MEG paradigm designed to induce visual gamma oscillations, adapting Hoogenboom’s approach [35]. Our visual stimulus was a circular sine wave grating, 5 degrees in diameter with a spatial frequency of 2 c.p.d., displayed at maximum contrast. We positioned this grating centrally on the screen, contracting towards a central fixation point. The grating’s initial speed was 2.2 degrees per second, increasing at random intervals between 50-3000 ms after stimulus onset. We instructed participants to press a button upon noticing any change in the grating’s contraction speed. After each response, we provided a 1000 ms rest period with feedback (“OK,” “early,” or “late”) to maintain engagement. Our experiment consisted of three runs, each comprising 80 trials. We maintained the projector’s refresh rate at 60 Hz for consistent visual presentation. In our analysis, we ignored the pseudo-random speed changes to isolate the neural correlates of gamma frequency oscillations, consistent with Hoogenboom’s method.

### 2.3 MEG recording

The study used a 275-channel CTF system with a 1200 Hz sampling rate, segmenting data into 4-second trials centered on stimulus onset. Synthetic Aperture Magnetometry (SAM) beamformer [58] analysis was employed to compare brain activity during 2-second baseline and post-stimulus periods within the 30-80 Hz gamma band. For the peak voxel identified in this SAM gamma-band image for each participant, the estimated time-series for this voxel was reconstructed for each trial.

### 2.4 Thalamo-Cortical Model (TCM)

We employed Dynamic Causal Modelling (DCM) [21] for steady-state responses to analyse spectral densities of neurophysiological signals. This approach utilises parameterised dynamical systems models; specifically state-space formulations of differential equation models (‘neural masses’), which describe the evolution of variables of interest (e.g. membrane potentials) as a function of connectivity parameters [21]. The thalamo-cortical (TCM) model incorporates interacting, layer-resolved cortical and thalamic populations, based on the architectures proposed by Douglas and Martin (2004) [19] and earlier work by Gilbert and Wiesel (1983) [24].

#### 2.4.1 TCM Overview

The thalamo-cortical model [66] is comprised of conductance-based equations of the forms of Hodgkin and Huxley and, subsequently, Morris and Lecar [48]. The model is comprised of pyramidal and interneuron populations in cortical layers 2/3 and 5, a stellate cell population in layer 4, a thalamic-projection pyramidal cell population in layer 6, and reticular and relay populations in the thalamus [79].

The dynamics of each population are determined by coupled differential equations:

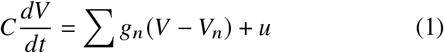

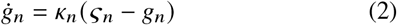

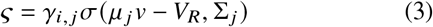

Here, *V* represents membrane potential, *g*_*n*_ denotes conductance change due to receptor *n, V*_*n*_ is the reversal potential of channel *n, C* is the membrane capacitance, and *u* includes external or internal input currents. *κ*_*n*_ is the decay rate for channel *n, γ*_*i, j*_ is the coupling parameter between populations *i* and *j*, and *σ* represents the sigmoid function for presynaptic depolarization.

Our model incorporates various conductance channels: AMPA, NMDA, GABA_*A*_, GABA_*B*_, M, and H (figure 1). M and H channels are exclusive to layer 6 thalamic-projection pyramidal cells and thalamic relay populations, which facilitate their characteristic bursting behavior. Our model is an extention of the NMDA model by Moran et al. (2011) [47] which includes a voltage-dependent mechanism for the NMDA magnesium block, with parameterised exponent a:

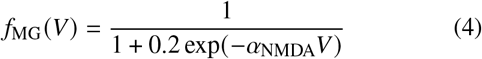

Reversal potentials and decay rates were adopted from established models and literature to enhance physiological accuracy, including specific parameters for GABA_*B*_, M-channels, and H-channels [23, 81, 63]. M-channels contribute to intrinsic cell membrane dynamics and excitation-inhibition balance, while H-channels influence resting membrane potential and may contribute to macroscopic network oscillations [12, 53, 60, 6]. GABA_*B*_ receptors have been linked to macroscale oscillations, particularly in the gamma range [13, 22].

**Figure 1:**
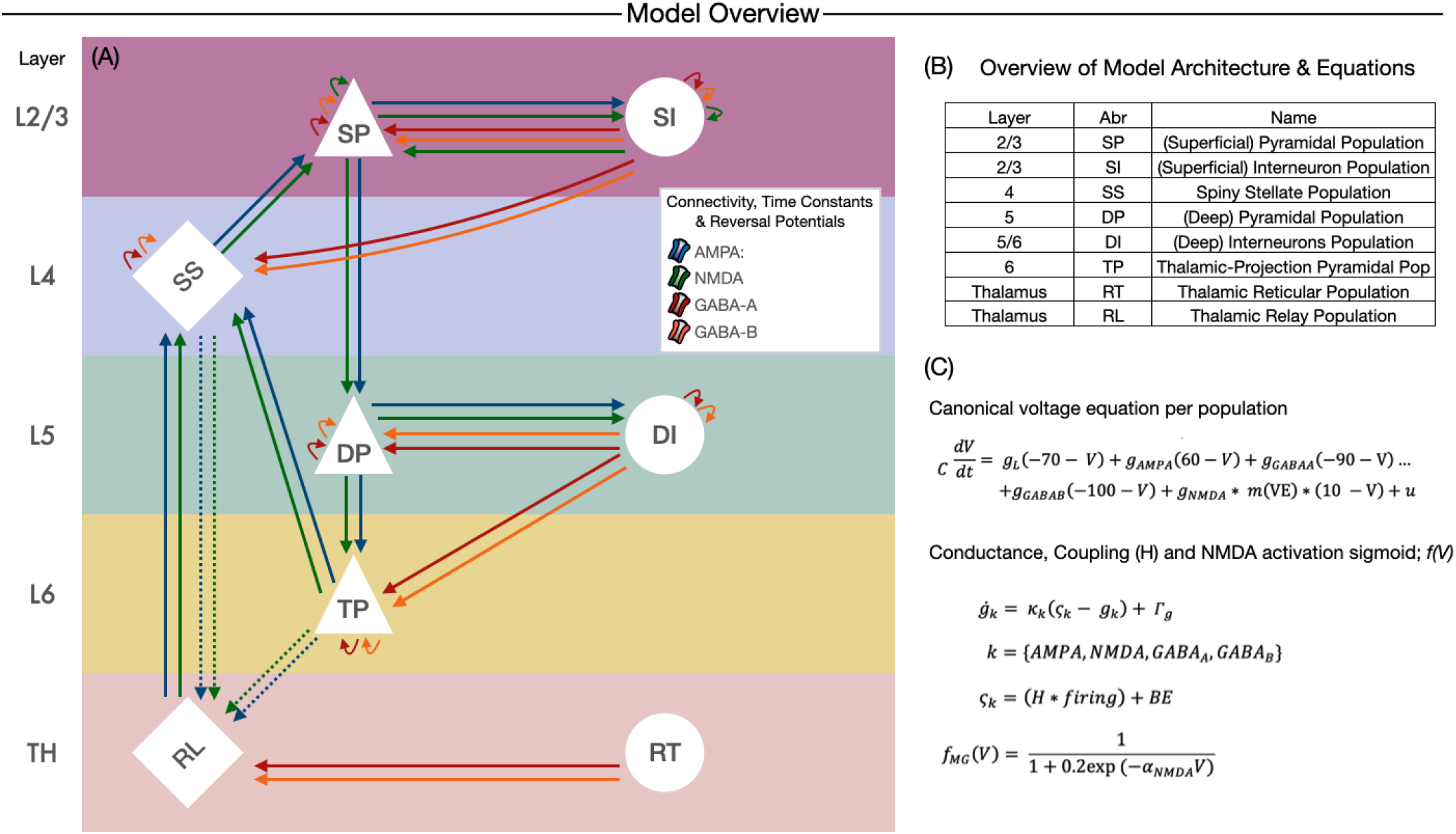
Thalamo-Cortical Model Architecture and Population Dynamics. **(A)** Schematic overview of the thalamo-cortical model architecture showing the connectivity between different neural populations across cortical layers (L2/3 to L6) and thalamic regions (TH). The model consists of six populations: Superficial Pyramidal (SP) and Interneuron (SI) in L2/3, Spiny Stellate (SS) in L4, Deep Pyramidal (DP) and Interneuron (DI) in L5, and Thalamic-Projection Pyramidal (TP) in L6. The thalamic populations include the Thalamic Relay (RL) and Thalamic Reticular (RT) populations. The model shows synaptic interactions mediated by AMPA (blue), NMDA (green), GABA-A (red), and GABA-B (orange) receptors, representing excitatory and inhibitory pathways. The connectivity lines are weighted according to time constants and reversal potentials. **(B)** Table summarizing the abbreviations for each population and their corresponding layers in the thalamo-cortical circuit. **(C)** The canonical voltage equation per population defines the dynamics of the membrane potential *V*, which includes contributions from leak conductance (*g*_*L*_), AMPA (2.2 ms delay), GABA-A (5 ms delay), GABA-B (300 ms delay), and NMDA (100 ms delay) receptor-mediated currents. Thalamic relay and layer 6 pyramidal populations further posses non-inactivating potassium channels (Kv7 or M-channels, 160 ms) and hyperpolarization-activated cation currents (H-channels, 100 ms) The nonlinear activation profiles of NMDA and H channels are modeled with sigmoid and inverse-signomid functions, respectively. NMDA conductance is modulated by voltage-dependent magnesium block represented by *f*_*MG*_ (*V*).

#### 2.4.2 Transfer Function

In accordance with other spectral DCM approaches, we employed the Laplace transform to compute the power spectrum of the model. The full procedure is outlined in the figure 2 panel “Generative model”. Briefly, we performed a local linearisation by numerically computing the Jacobian of the system about a fixed point. From this linearised approximation of the model, the Laplace transform at frequency i, in hertz, is given by:

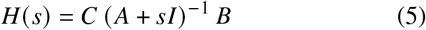

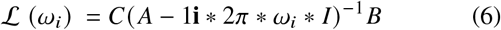

Here (5) denotes the “canonical form” Laplace transform of a system, where *H(s)* is the transfer function and *s* is the complex Laplace variable. Equation (6) represents the Laplace transform evaluated at a specific frequency, *w(i)*, in hertz. A is the system Jacobian matrix (*df/dx*), B the input Jacobian (*df/du*) and C a vector of (fixed) weights controlling the contribution of each state element to the output local field potential. This transfer function approach is computationally efficient compared with numerically integrating the model equations. In contrast to conventional spectral DCM, we did not incorporate any spectral noise components explicitly, such as 1/f “aperiodic” components or a discrete cosine basis set to represent neuronal fluctuations in the frequency domain. Thus, the model output comprises only the Laplace transform of the model fitted to data and under the assumption of a white noise input.

**Figure 2:**
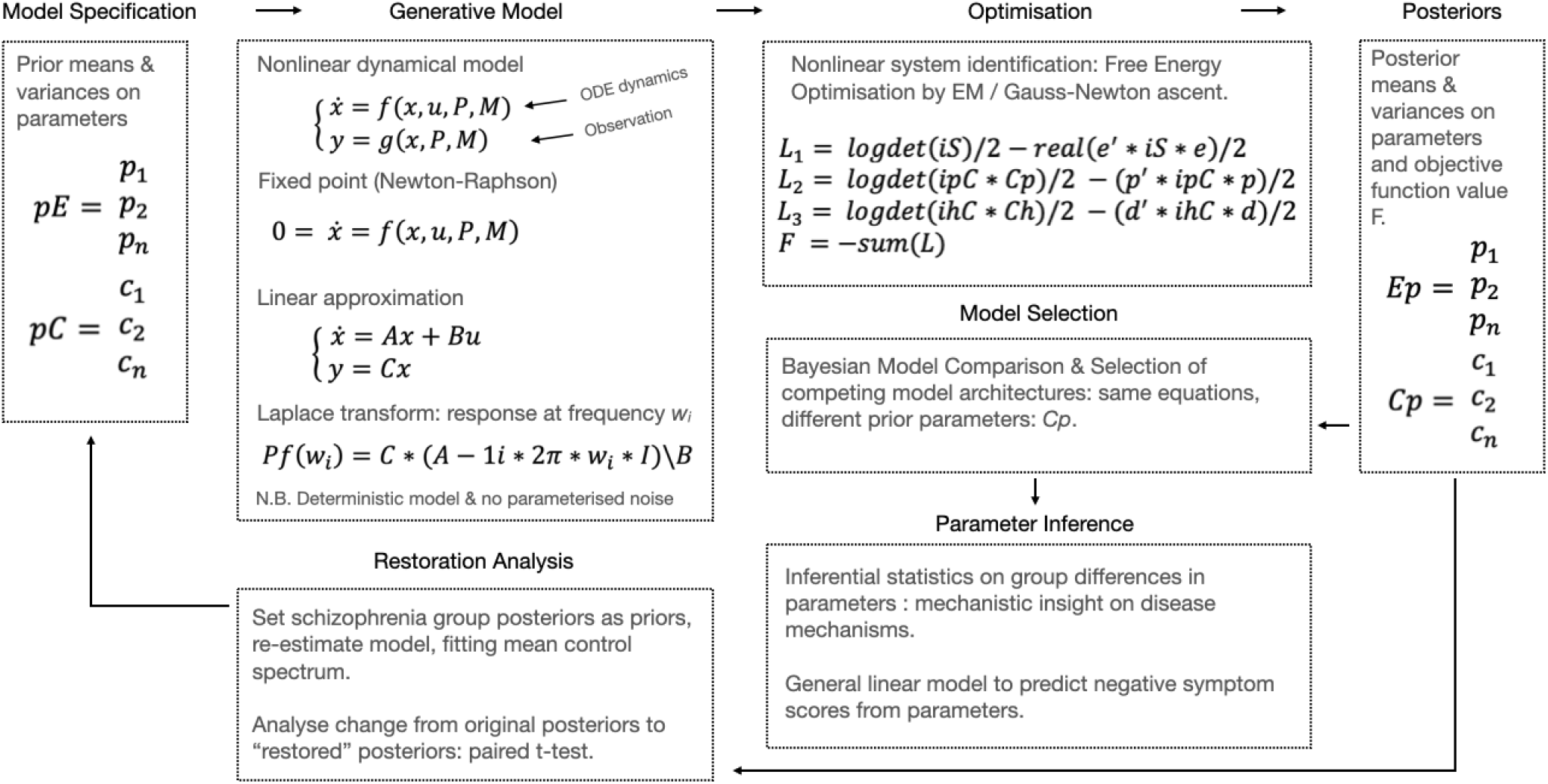
Schematic representation of the generative model pipeline for model optimization and parameter inference. The process begins with **Model Specification**, where prior means (*pE*) and variances (*pC*) on the parameters are defined. The **Generative Model** uses a nonlinear dynamical system represented by ordinary differential equations (ODEs), and applies fixed point and linear approximations to solve the system, including Laplace transform for response at frequency *ω*_*i*_. In the **Optimisation** stage, the system undergoes parameter estimation using free energy optimization via Expectation Maximization (EM) or Gauss-Newton ascent. The resulting posterior means (*E p*) and variances (*C p*) on the parameters are used for **Model Selection**, where Bayesian model comparison selects the best fitting model architecture. **Parameter Inference** involves statistical analysis on group differences in parameter estimates, providing mechanistic insights into disease mechanisms. In a final step, **Restoration Analysis** we re-estimate parameters using schizophrenia group posteriors and performs t-tests to analyze differences with the control group.

The TCM implementation contains 56 variables consisting of 8 populations, with 7 states each: mV, *g*_AMPA_, *g*_GABA-A_, *g*_NMDA_, *g*_GABA-B_, *g*_*M*_, and *g*_*H*_

The cortex-thalamus-cortex feedback loop in the TCM incorporates a parameterized delay, initially set to 11 ms (comprising 8 ms from cortex to thalamus and 3 ms from thalamus to cortex). This delay was subject to variation during model fitting to accommodate discrepancies in literature-based estimates [42, 1, 32].

### 2.5 Bayesian Model Comparison (BMC)

To evaluate the relative explanatory power of different model configurations within our thalamo-cortical framework, we employed Bayesian model comparison [75, 56]. This approach allows us to quantitatively assess which model structure best explains the observed MEG data, balancing model complexity against goodness of fit.

We constructed a model space comprising 15 variations of the TCM, each representing different hypotheses about thalamocortical interactions (fig 3). The first 4 models contained connectivity mediated only by AMPA, NMDA, GABA_*A*_ or GABA_*B*_. The next 6 models tested every coupling of these 4 receptor systems, followed by a further 4 models testing every trio, and the final model featured all four receptor systems. That is, models 1:14 were all nested models within model 15.

**Figure 3:**
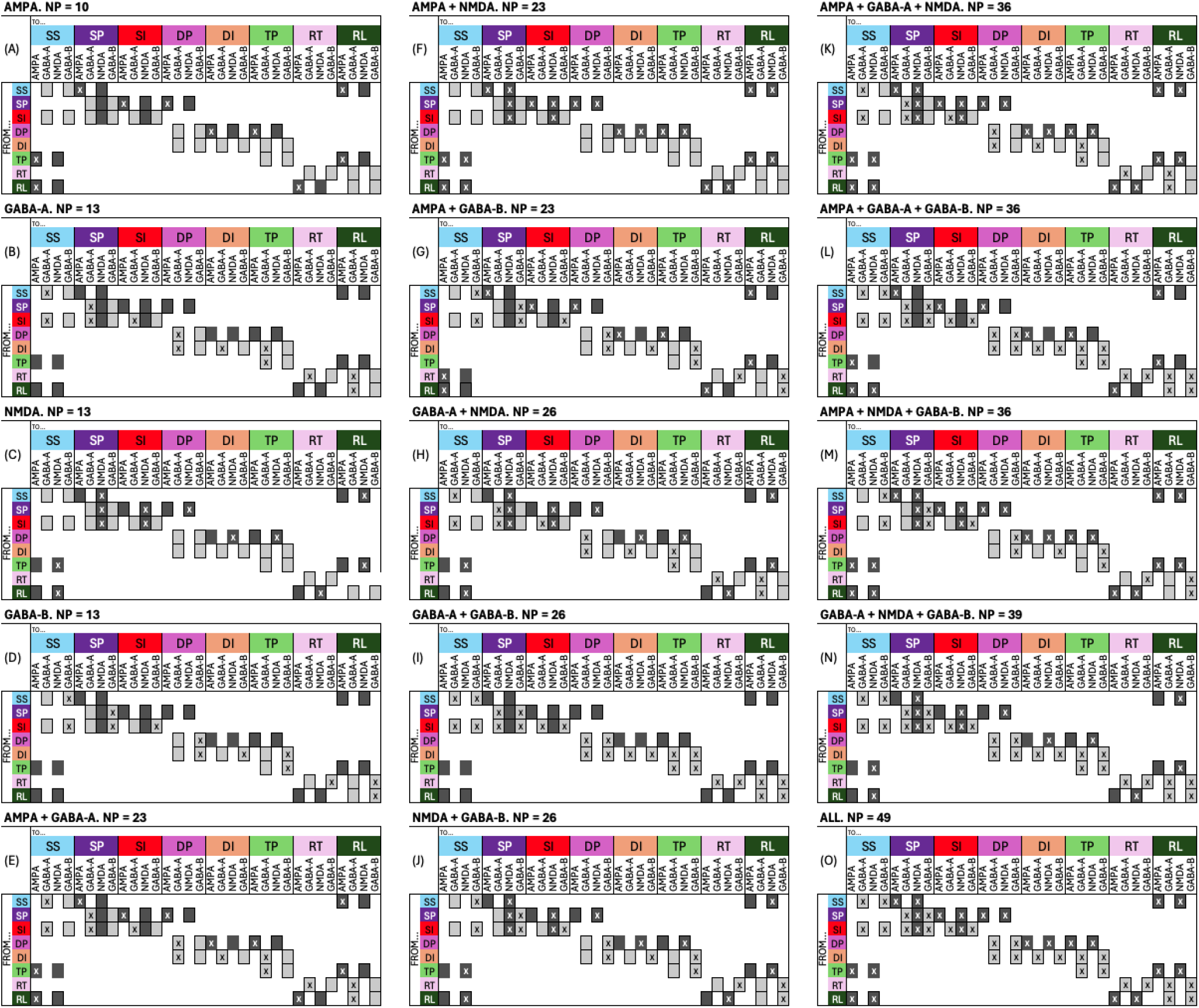
Connectivity matrices across different synaptic combinations. Each panel (A-O) represents the connectivity structure for a specific synaptic receptor combination: AMPA (**A**), GABA-A (**B**), NMDA (**C**), GABA-B (**D**), AMPA + GABA-A (**E**), AMPA + NMDA (**F**), AMPA + GABA-B (**G**), GABA-A + NMDA (**H**), GABA-A + GABA-B (**I**), NMDA + GABA-B (**J**), AMPA + GABA-A + NMDA (**K**), AMPA + GABA-A + GABA-B (**L**), AMPA + NMDA + GABA-B (**M**), GABA-A + NMDA + GABA-B (**N**), and all synaptic types (**O**). Each matrix illustrates the connections between neuronal populations (SP, SI, SS, DP, DI, TP, RL, RT) for each synaptic mechanism. Crosses within the squares denote existing connections that were allowed to vary during optimization whereas the absence of a cross denotes fixed connections during optimization. These submodels were subsequently used in the Bayesian Model Comparison.

For each model, we computed the log-evidence, which quantifies the trade-off between model accuracy and complexity. This metric is derived via the free energy approximation used in the DCM optimization process [51]. Models with higher log-evidence are considered to provide a better explanation of the data.

We then used both fixed effects (FFX) and random effects (RFX) Bayesian model selection to account for potential inter-subject variability in the optimal model structure. The FFX analysis assumes all subjects’ data are best explained by the same model, while RFX allows for different optimal models across subjects.

For the RFX analysis, we calculated several metrics. The expectation of the posterior represents the average or expected value of the model frequencies across the population. Exceedance probabilities indicate the likelihood that a particular model is more frequent in the population than all other considered models. We also computed protected exceedance probabilities, which provide a more conservative estimate of model superiority. These consider the possibility that observed differences in model evidence could be due to chance, thus offering protection against false positives when comparing models.

Collectively, these metrics help us assess which models are most likely to explain the observed data across the population, while accounting for individual variability and the possibility of chance findings.

Additionally, we computed the Bayesian Omnibus Risk (BOR) to quantify the probability that observed model differences are due to chance [56]. We performed this analysis for three dataset groupings: all datasets combined, schizophrenia datasets, and control datasets.

### 2.6 Parameter Inference

We applied the TCM to individual participant data, generating a set of parameter estimates for each subject. To assess group differences between healthy controls and individuals with established schizophrenia, we conducted independent t-tests for each model parameter.

To address the issue of multiple comparisons arising from testing multiple model parameters, we employed the False Discovery Rate (FDR) correction method. The FDR correction was applied to the p-values obtained from the t-tests.

### 2.7 Principle Component Analysis (PCA)

In our previous study, where we compared visual gamma activity between individuals with established schizophrenia and healthy controls, we demonstrated that we could distinguish between these two groups based on their MEG signals [67]. For the current study, we incorporated the model parameters from the TCM to investigate whether we could achieve better group differentiation. We conducted Principal Component Analysis (PCA) on the MEG power spectrum and the model parameters to explore this potential improvement in group separation.

### 2.8 Negative Symptoms

Our study extended beyond identifying group differences in model parameters to examining their relationship with clinical symptoms. Without strong prior hypotheses regarding neurobiological mechanisms underwriting negative symptoms - and acknowledging the “multi-system” nature of psychiatric physiology, we employed data-driven, stepwise regression to determine which parameters were significantly correlated to negative symptoms in individuals with schizophrenia. The stepwise regression systematically evaluated combinations of model parameters, iteratively including or excluding them based on their association strength with negative symptom severity. This data-driven approach allows for an exploration of the links between neurophysiological parameters and symptom severity.

### 2.9 Parameter Restoration Analysis

To further investigate the differences between healthy controls and individuals with schizophrenia, we performed a ‘restoration’ analysis. This approach quantifies the extent to which parameters from the schizophrenia group need to be adjusted to match the spectral characteristics observed in the control group.

Initially, we set the posterior parameter estimates obtained from the schizophrenia group as prior distributions for a new model estimation. Utilizing these priors, we re-estimated the TCM by fitting it to the mean spectrum of the control group, resulting in a set of “restored” posterior parameters. Like the original model fits, this re-optimisation also employed the Variational Laplace routine in DCM, which optimises a free energy objective function. Therefore, this restoration analysis assumes that the movement in synaptic connectivity necessary to move the neurophysiological signature of an individual with schizophrenia to that of the mean control individual, follows the path described by minimising free energy.

We analyzed the changes from the original schizophrenia group posteriors to these restored posteriors to understand the parametric differences that account for the spectral disparities between the two groups. To quantify the significance of these changes, we conducted paired t-tests for each parameter.

This analysis provides a quantitative measure of the neurophysiological differences between the two groups in terms of TCM parameters, highlighting potential areas of dysfunction in schizophrenia that could serve as potential therapeutic targets.

## 3 Results

### 3.1 Spectral Power Responses

Analysis of the broadband power spectrum revealed significant differences between the schizophrenia group and healthy controls (Figure 4A) [F(1, 29) = 4.15, P = .047] with the SZ group showing an estimated mean reduction of 3 Hz (Controls mean = 58, SE = 0.92, SZ mean = 55, SE = 0.95). Most notably, the schizophrenia group exhibited an increased alpha amplitude compared to the control group, with a prominent peak observed in the 8-12 Hz frequency range. This elevation in alpha power was particularly pronounced around 10 Hz.

**Figure 4:**
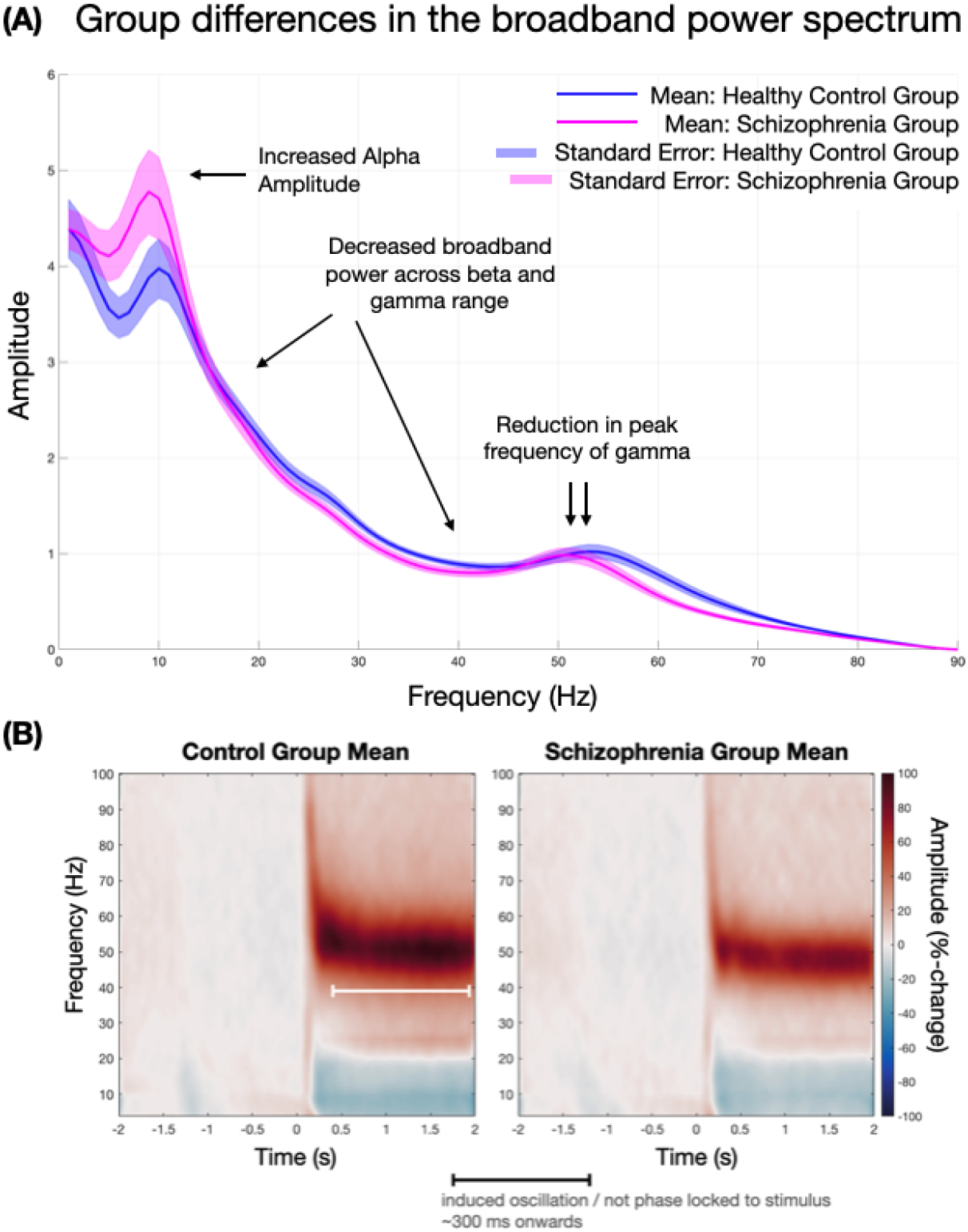
Comparison of brain activity patterns between healthy controls and individuals with schizophrenia. **(A)** Group differences in the broadband power spectrum. The graph shows mean amplitude across frequencies for healthy controls (blue) and schizophrenia patients (pink), with standard error bands. Key differences include increased alpha amplitude, decreased broadband power across beta and gamma ranges, and reduced peak frequency of gamma in the schizophrenia group.**(B)** Time-frequency representations of mean brain activity for control and schizophrenia groups. The heatmaps display frequency (y-axis) over time (x-axis), with color intensity representing amplitude changes. An induced oscillation period is noted, occurring approximately 300 ms onwards and not phase-locked to the stimulus. The schizophrenia group shows altered patterns of activity, particularly in higher frequency ranges, compared to controls.

In contrast to the alpha band, the schizophrenia group showed decreased broadband power across the beta (13-30 Hz) and gamma (30-80 Hz) ranges relative to healthy controls. This reduction in high-frequency power was accompanied by a subtle shift in the peak frequency of gamma oscillations, with the schizophrenia group displaying a lower peak frequency compared to controls.

While the power spectrum showed increased alpha amplitude in the schizophrenia group, the time-frequency plots revealed this as a more pronounced alpha desynchronization (power decrease relative to baseline) following stimulus presentation. This apparent discrepancy is due to the different nature of these analyses, with power spectra representing absolute power and time-frequency plots showing relative changes.

Additionally, the time-frequency analysis highlighted reduced induced high-frequency oscillations in the schizophrenia group, particularly in the gamma range, consistent with the decreased broadband power observed in the spectrum analysis.

Detailed information about the spectrum analysis can be found in [67] where these findings were reported originally.

### 3.2 Bayesian Model Comparison Analysis

The Bayesian Model Comparison (BMC) analysis evaluated 15 variants of the thalamo-cortical model across all datasets, as well as separately for schizophrenia and control groups. FFX analysis consistently showed high posterior probabilities for the most comprehensive model, which included all possible connections. This pattern was observed across all dataset groupings (figure 5). RFX analysis, accounting for potential inter-subject variability, provided supporting evidence through the expectation of the posterior, exceedance probabilities, and protected exceedance probabilities. These metrics also favored the fully connected model. BOR values were calculated for all dataset groupings, resulting in extremely low values. This indicates strong evidence that the observed preferences for the fully connected model were not due to chance.

**Figure 5:**
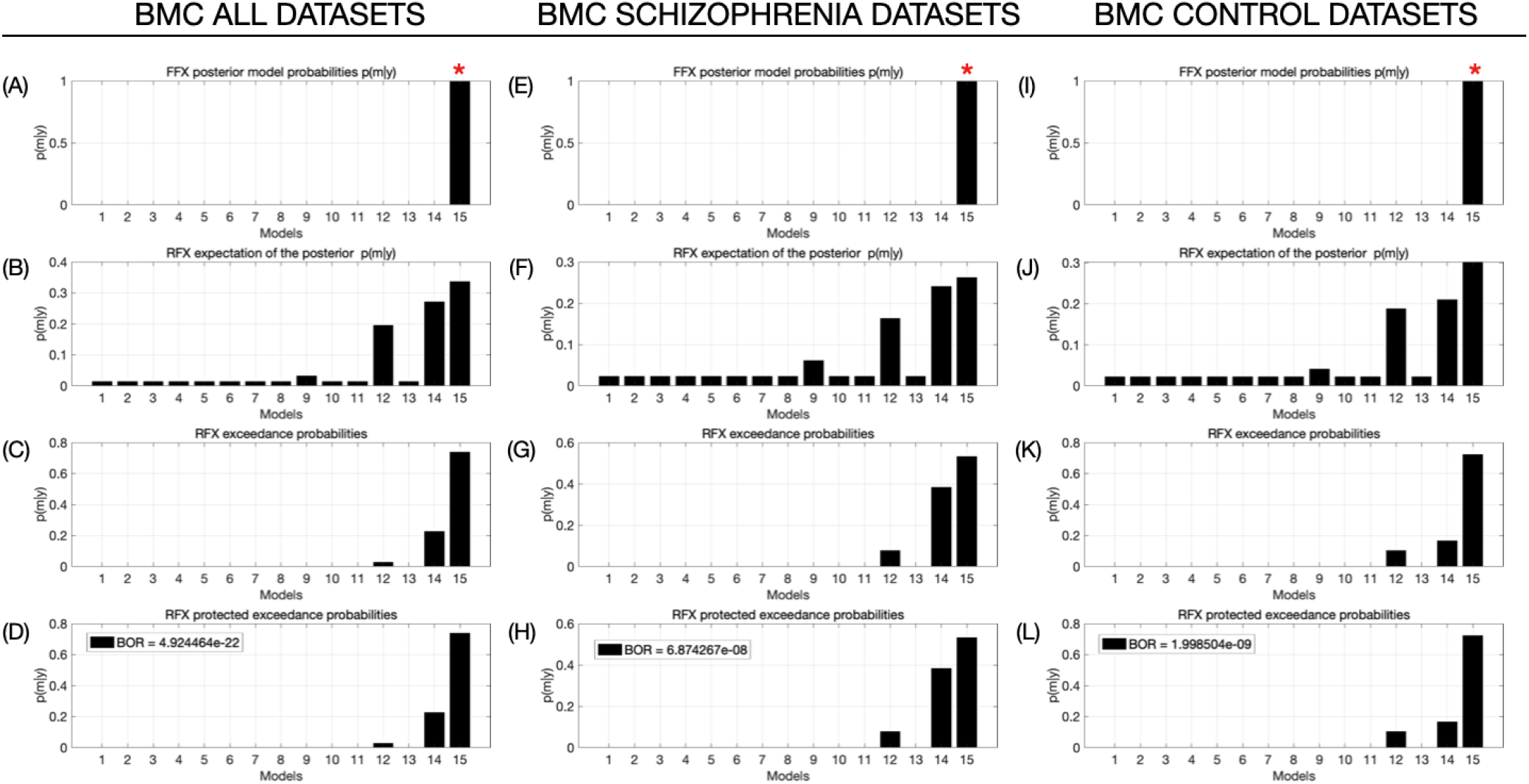
Fixed Effects (FFX) and Random Effects (RFX) Bayesian Model Comparison (BMC) analysis. The figure is organized into three main columns representing different dataset groupings: all datasets combined, schizophrenia datasets, and control datasets. Each column contains four rows of graphs (A-D, E-H, I-L) showing different aspects of the BMC analysis: **(A,E,I)** FFX posterior model probabilities: These graphs show the fixed effects analysis results, with model 15 consistently emerging as the winning model (marked with a red asterisk) across all dataset groupings. **(B,F,J)** RFX expectation of the posterior: These display the random effects analysis, showing the expected posterior probability for each model. Models 14-15 generally show higher probabilities, particularly in the control datasets. **(C,G,K)** RFX exceedance probabilities: These graphs indicate the probability that each model exceeds all others in the random effects analysis. Model 15 consistently shows the highest exceedance probability across all dataset groupings. **(D,H,L)** RFX protected exceedance probabilities: Similar to the exceedance probabilities, but protected against the possibility of all models being equally likely. The Bayes Omnibus Risk (BOR) values are provided, showing extremely low probabilities that all models are equally likely.This figure illustrates the results of both fixed and random effects analyses, consistently favoring model 15 across different analytical approaches and dataset groupings.

**Figure 6:**
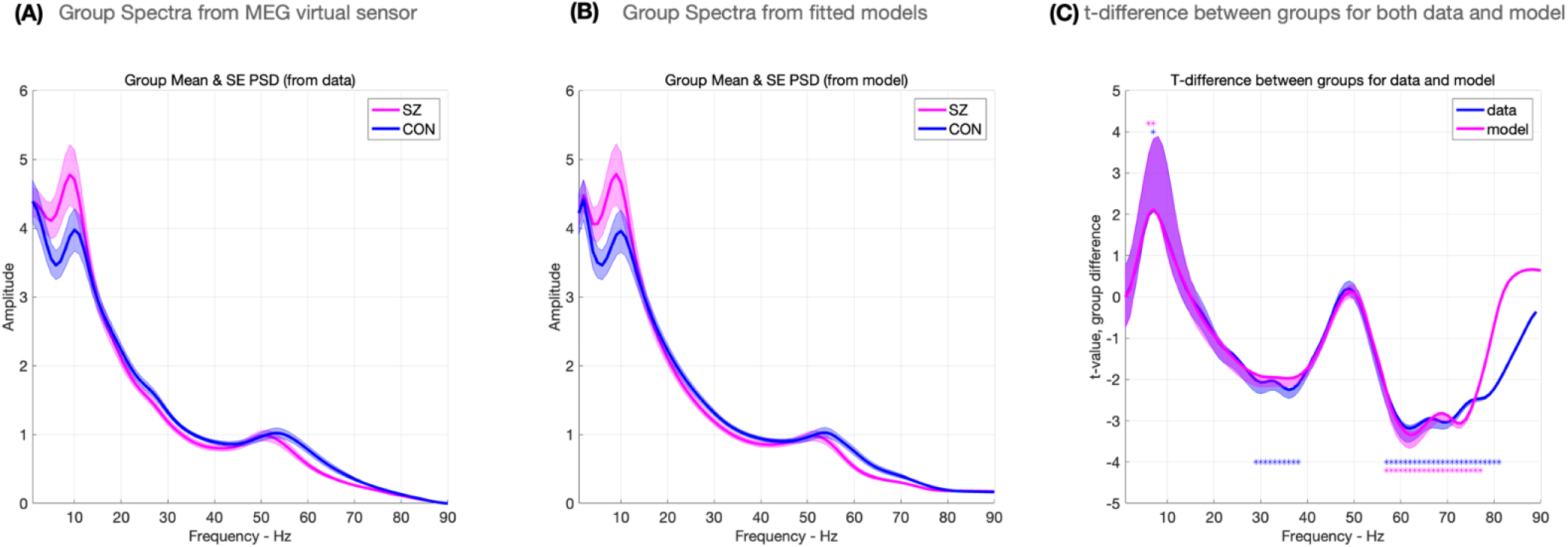
Comparison of empirical MEG data and model-fitted spectra. **(A)** Group Spectra from MEG virtual sensor: This graph shows the group mean and standard error of the power spectral density (PSD) derived directly from MEG data for both SZ (pink) and CON (blue) groups across frequencies from 0 to 90 Hz. **(B)** Group Spectra from fitted models: This graph displays the group mean and standard error of the PSD as predicted by the fitted computational models for both SZ and CON groups. The close resemblance to panel A indicates a good model fit for both groups. **(C)** t-difference between groups for both data and model: This graph illustrates the t-statistic of the difference between SZ and CON groups across frequencies, comparing the empirical data (blue) and model predictions (pink). The similarity between the two lines demonstrates how well the model captures the group differences observed in the actual data. Overall, this figure demonstrates the high quality of the model fit for both groups. The model successfully captures the spectral characteristics and group differences observed in the empirical MEG data, validating its use for further analysis of spectral alterations in schizophrenia.

Thus, the BMC results suggest that a complex, fully connected thalamo-cortical circuit best explains the observed MEG data in both healthy controls and individuals with schizophrenia (figure 5).

### 3.3 Parameter Comparison

The parameter comparison between the control and schizophrenia groups revealed significant differences in specific neural connection strengths within the thalamo-cortical model.

Notably, the NMDA-mediated self-connection within superficial pyramidal (SP) cells showed a significant increase in the schizophrenia group compared to controls:

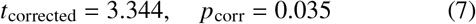

This suggests an enhancement of recurrent excitation within superficial cortical layers in individuals with schizophrenia.

Conversely, the GABA-B-mediated connection from superficial interneurons (SI) to SP cells was significantly reduced in the schizophrenia group:

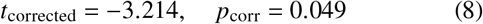

This finding indicates a potential deficit in slow inhibitory control within the superficial cortical layers in schizophrenia.

### 3.4 Principal Component Analysis (PCA)

PCA was applied to both the power spectra and model parameters to assess group separation between healthy controls and individuals with schizophrenia. The results reveal a striking differentiation between the two groups when visualized in a two-dimensional space (Figure 8). The first principal component derived from the model parameters is plotted against the principal component of the power spectra. This representation demonstrates a clear separation between the control group and the schizophrenia group, with minimal overlap between the two clusters.

**Figure 7:**
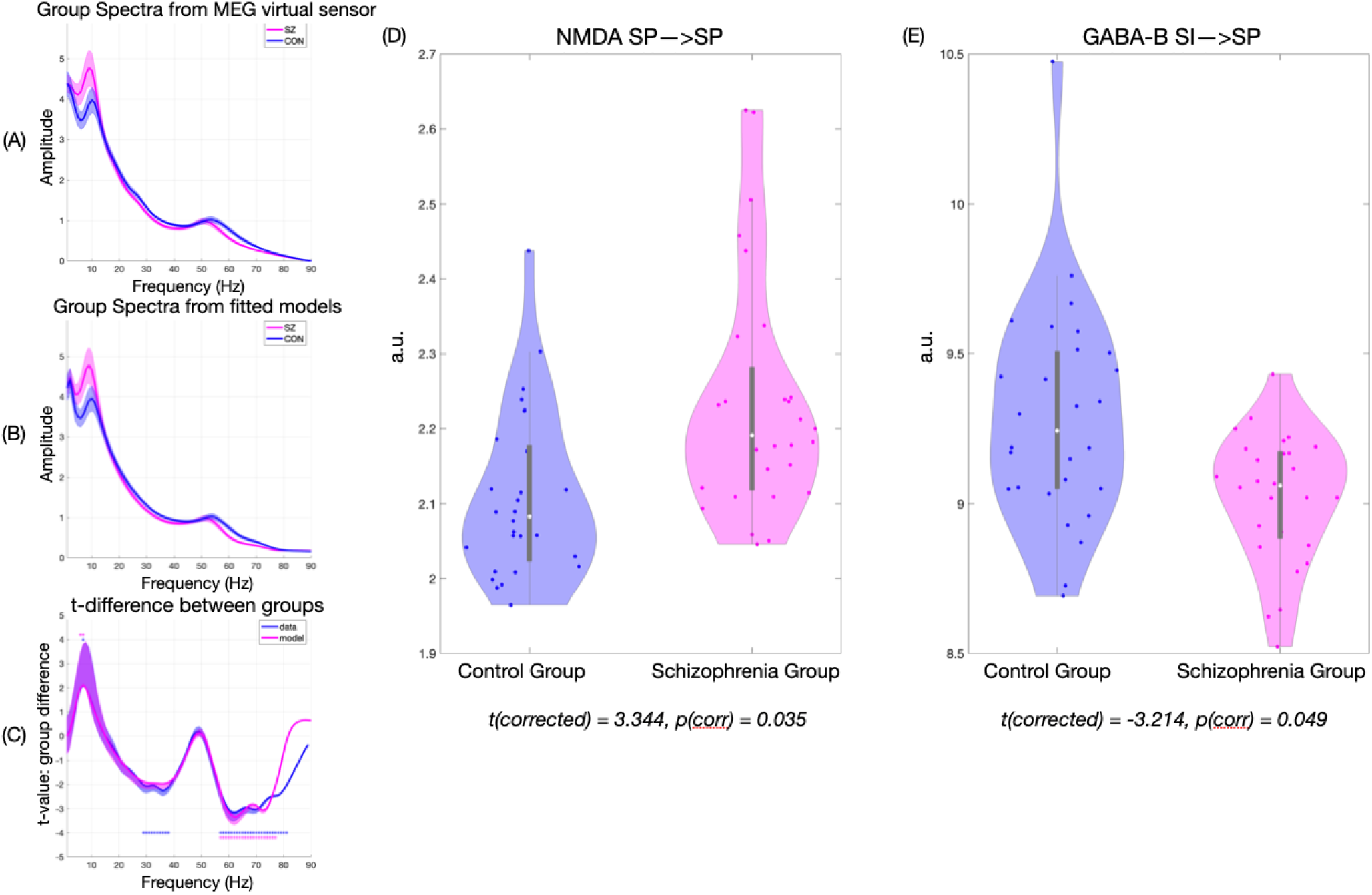
Spectral and synaptic parameter comparison between control and schizophrenia groups. Panels **A** and **B** show power spectra from MEG data and fitted models respectively, with amplitude plotted against frequency. Panel **C** displays the t-difference between groups for both data and model. Panels **D** and **E** are violin plots illustrating group differences in specific synaptic parameters: NMDA SP→SP connectivity **(D)** shows higher values for the schizophrenia group, while GABA-B SI→SP connectivity **(E)** shows lower values. These are both statistically significant using an independent t-test (FDR corrected)

**Figure 8:**
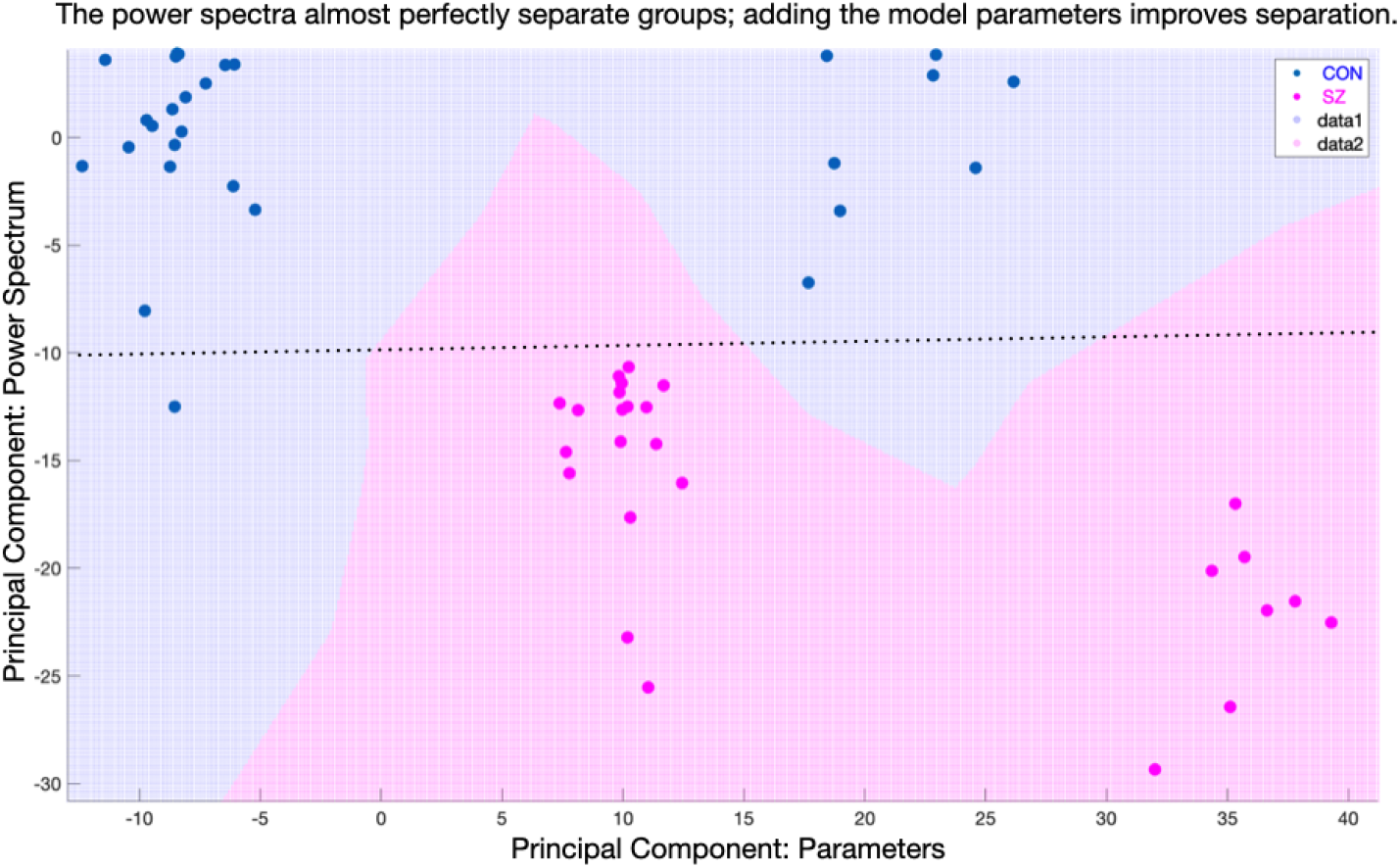
Principal Component Analysis (PCA) of power spectra and model parameters. The x-axis represents the principal component derived from model parameters, while the y-axis shows the principal component from power spectrum data. Blue dots indicate control subjects, and pink dots represent individuals with schizophrenia. The plot demonstrates clear separation between the two groups.

Although power spectra alone offer significant group separation, incorporating model parameters further enhances this distinction, revealing two subgroups within both the control and schizophrenia groups.

### 3.5 Negative Symptoms

A stepwise regression analysis using a Generalized Linear Model (GLM) was conducted to identify parameters predictive of negative symptoms in schizophrenia. The final model achieved statistical significance (*F* = 7.74, *p* = 0.001) and explained a substantial portion of the variance in negative symptom scores (*r*^2^ = 0.51, adjusted *r*^2^ = 0.45).

The model identified three key parameters as predictors of negative symptom severity: AMPA-mediated connectivity from thalamic projection (TP) neurons to reticular (RL) neurons (AMPA TP→RL), GABA-B-mediated self-inhibition of superficial interneurons (GABA_B_ SI→SI), and an interaction term between AMPA TP→RL and GABA_B_ SI→SI. The analysis revealed a positive correlation between predicted and observed negative symptom scores, indicating that the model generally captures the trend in symptom severity. The AMPA TP→RL parameter showed a strong negative association with symptom severity, while the GABA_B_ SI→SI parameter demonstrated a positive association. The interaction term between these two parameters showed a positive relationship with symptom severity.

These findings suggest that alterations in thalamocortical AMPA signaling and local GABA-B-mediated inhibition, as well as their interaction, may play significant roles in the manifestation of negative symptoms in schizophrenia.

### 3.6 Parameter Restoration

Our parameter restoration analysis demonstrated significant changes across multiple synaptic connections in the ThalamoCortical TCM when adjusting the schizophrenia group parameters to match the spectral characteristics of the control group (figure 10).

**Figure 9:**
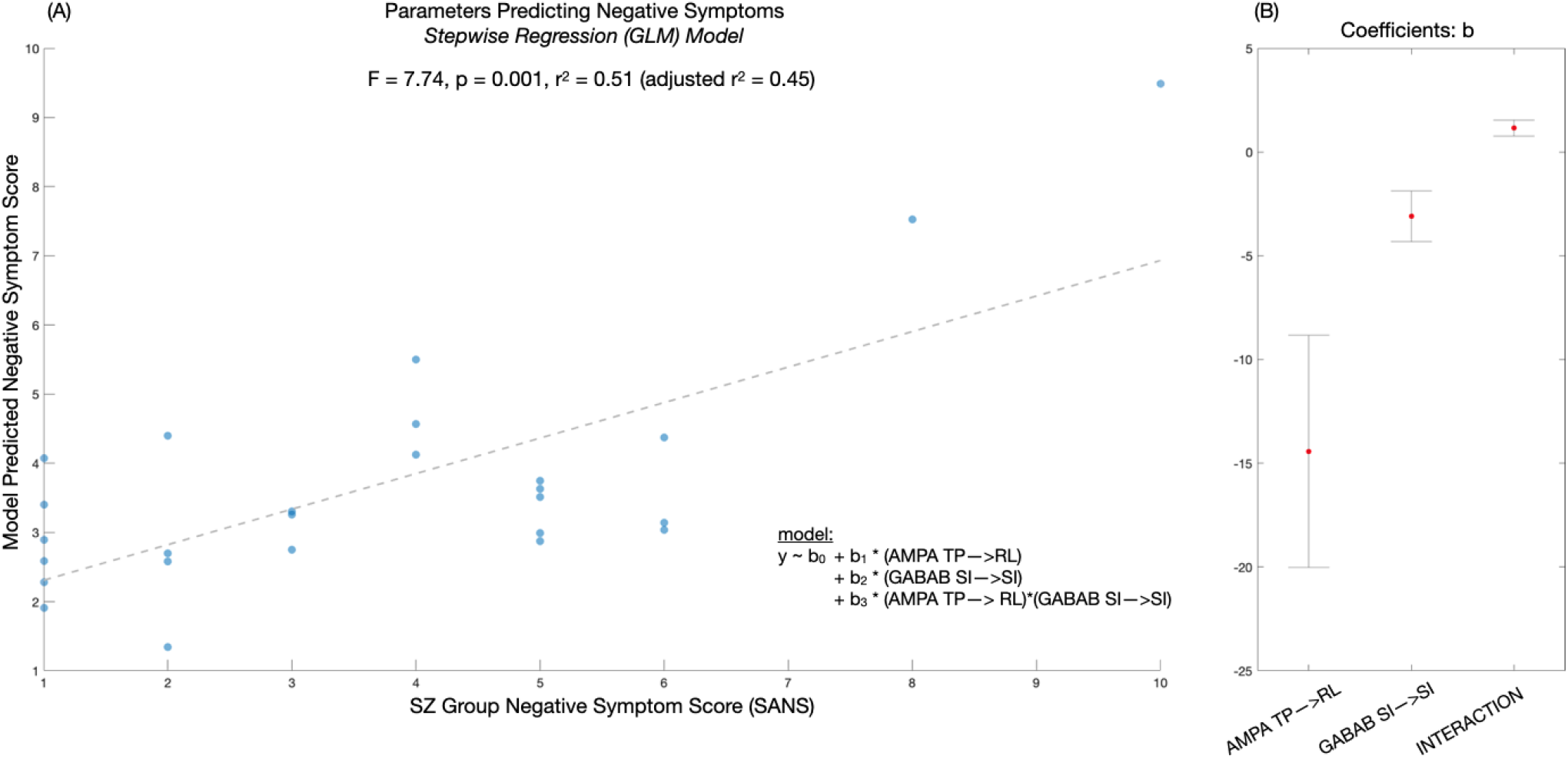
Stepwise regression model predicting negative symptoms in schizophrenia. **(A)** Scatter plot comparing observed negative symptom scores (SANS) with predicted scores from the stepwise regression model. Each point represents an individual with schizophrenia. The dashed line shows the model fit (F = 7.74, p = 0.001, r^2^ = 0.51, adjusted r^2^ = 0.45). The model equation includes synaptic parameters: AMPA (TP→RL), GABAB (SI→SI), and their interaction. **(B)** Coefficient plot showing the relative contributions (beta weights) of each predictor in the model. Error bars indicate 95% confidence intervals.

**Figure 10:**
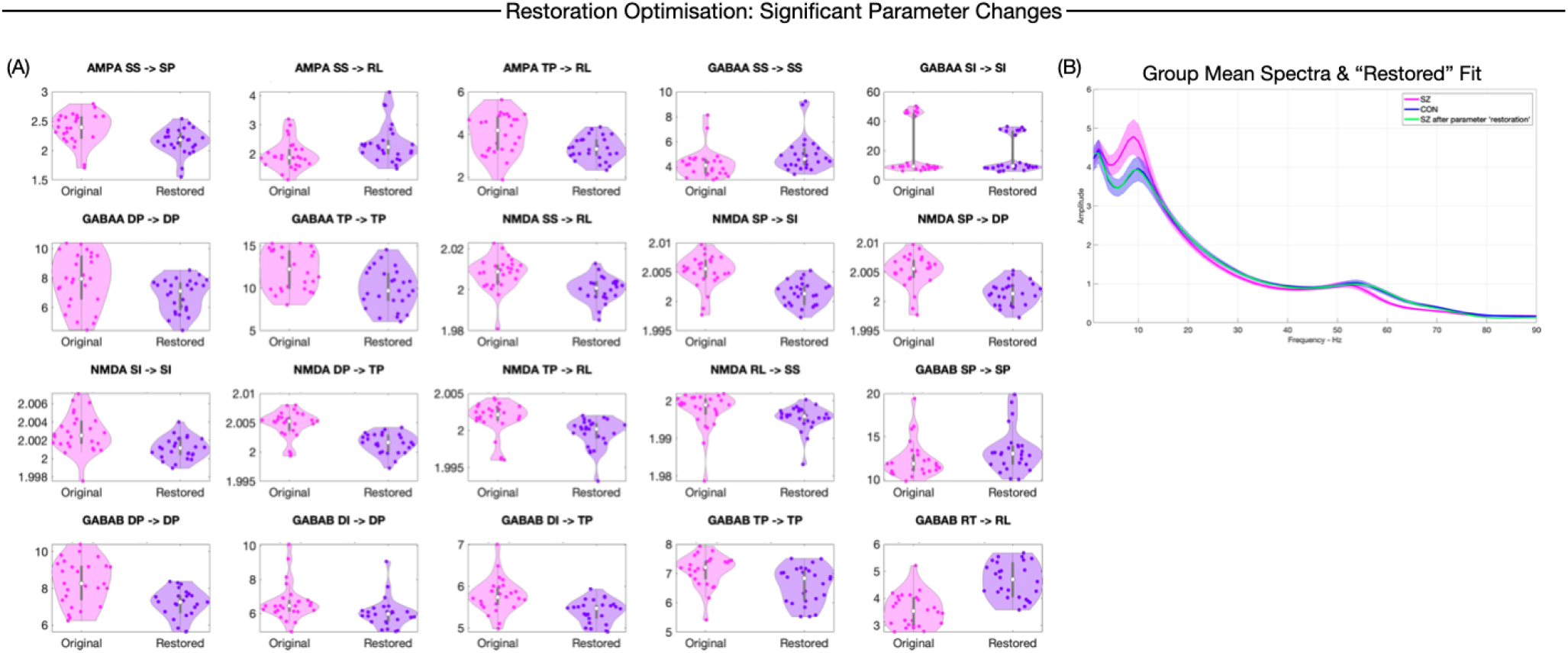
Restoration Optimisation: Significant Parameter Changes and Spectral Effects. **(A)** Multiple violin plots showing changes in synaptic parameters between the original schizophrenia group (pink) and the computationally “restored” state (purple). Each plot represents a different synaptic connection. Most importantly, more parameters need to be adjusted significantly (FDR corrected) to match the power spectrum of the healthy controls than we found differed significantly between schizophrenia and healthy controls in our initial analysis. **(B)** Power spectra comparison showing the original control group (blue), schizophrenia group (pink), and the “restored” schizophrenia model (green). The x-axis represents frequency (Hz), and the y-axis represents amplitude.

Key findings from the paired t-tests comparing original schizophrenia posteriors to restored posteriors include significant alterations in AMPA-mediated, GABA-A-mediated, NMDA-mediated, and GABA-B-mediated connections. Specifically, AMPA-mediated connections from Superficial Stellate (SS) to Superficial Pyramidal (SP) and SS to Reticular (RL) exhibited significant decreases after restoration, as did the Thalamic Projection (TP) to RL connection. In GABA-A-mediated responses, SS self-inhibition increased substantially, and Superficial Interneuron (SI) self-inhibition also showed a significant increase, while Deep Pyramidal (DP) and TP self-inhibitions decreased significantly. For NMDA-mediated connections, most such as SS to RL, SP to SI, SP to DP, SI to SI, DP to TP, and TP to RL displayed slight but significant decreases, with the RL to SS connection marginally but significantly increasing. Lastly, GABA-B-mediated interactions saw a significant increase in SP self-inhibition, significant decreases in DP self-inhibition, DI to DP, DI to TP, and TP self-inhibition, and a substantial increase in the Reticular Thalamus (RT) to RL connection. These parameter adjustments resulted in a model that produced spectral outputs more closely aligned with the control group, particularly in the alpha and gamma frequency ranges. These results quantify the extent of neurophysiological differences between schizophrenia patients and healthy controls in terms of TCM parameters. The significant changes across multiple neurotransmitter systems and synaptic connections highlight the complex nature of the neural dysregulation in schizophrenia. The parameters requiring the most substantial adjustments, such as GABA-A mediated SS self-inhibition and GABA-B mediated RT to RL connection, may represent key areas of dysfunction in schizophrenia and could serve as potential therapeutic targets for future investigations.

## 4 Discussion

In this study, we employed a thalamo-cortical conductance-based model to investigate neural circuit dynamics in schizophrenia using MEG data. Our approach combined spectral analysis, Dynamic Causal Modelling, and parameter restoration.

### 4.1 Spectral Analysis

Our findings reveal distinct alterations in neural oscillatory patterns in schizophrenia. The most prominent observation is the significantly increased alpha (8-12 Hz) power in the schizophrenia group relative to healthy controls. This may suggest an impairment in alpha modulation, potentially related to deficits in attentional control and sensory gating that are frequently observed in schizophrenia [80]. In contrast to the heightened alpha power, we observed decreased power in the beta (13-30 Hz) and gamma (30-80 Hz) frequency ranges in the schizophrenia group. This finding is consistent with previous research [64, 67, 31, 11], which provided robust evidence for reduced gamma-band activity across various paradigms in schizophrenia. Our results extend this finding to the beta range, suggesting broader deficits in high-frequency oscillations.

A more subtle but noteworthy observation is the reduction in peak gamma frequency in the schizophrenia group. While altered gamma synchrony in schizophrenia has been extensively studied [73, 80], changes in peak gamma frequency have also received significant attention[29, 77, 27], providing additional insights into neural oscillation abnormalities in the disorder. This shift in frequency reflects an imbalance in excitatory-inhibitory dynamics within cortical microcircuits [26, 66].

Collectively, these oscillatory alterations suggest disrupted neural dynamics in schizophrenia, characterized by an imbalance between low- and high-frequency oscillations. Crucially, it is only by employing computational modelling across these broad-band spectral group differences, that we are able to demonstrate the thalamo-cortical circuit mechanisms related to schizophrenia pathophysiology and negative symptom severity.

### 4.2 Parameter Inference and Excitation/Inhibition Imbalance

In the parameter inference we identified key changes in synaptic connectivity in schizophrenia. Specifically, we observed an increase in NMDA-mediated recurrent excitation among superficial pyramidal cells (*t*_corrected_ = 3.344, *p*_corr_ = 0.035) and a decrease in GABA-B-mediated inhibition from superficial interneurons to superficial pyramidal cells (*t*_corrected_ = ™3.214, *p*_corr_ = 0.049). This finding replicates and extends our previous results using a simpler model on the same dataset, which also identified alterations in SP->SP self-connections in schizophrenia [67].

Crucially, the parameter changes observed in this study reconcile the excitatory/inhibitory (E/I) balance theory in schizophrenia (that integrates both glutamatergic and GABAergic dysfunction) [15, 72] with the NMDA-mediated “dysconnectivity” hypothesis. The E/I imbalance hypothesis assumes that the ratio of excitatory to inhibitory neurotransmission is disrupted in schizophrenia, leading to aberrant neural circuit function and, consequently, to the diverse symptoms of the disorder [38].

The increased NMDA-mediated excitation we observed in our sample of individuals with established schizophrenia is particularly interesting, as it appears to contradict previous findings in schizophrenia that showed loss of pyramidal synaptic gain [2] and the NMDA receptor hypofunction hypothesis of schizophrenia [45, 49, 14]. This hypothesis states that NMDA receptor function is reduced in chronic schizophrenia, contributing to various cognitive and behavioral symptoms. Our finding of increased NMDA-mediated excitation in this population is unexpected and challenges previous results. Several interpretations of this result are possible:

#### 1. Hyperfunction/Hypofunction

Our results bear a close resemblance to the hyperglutamatergic state often observed in first-episode schizophrenia or individuals at clinical high-risk of psychosis. Early stages of the disorder are characterized by NMDA receptor hyperfunction, which later transitions to hypofunction, as a compensatory mechanism, in chronic stages [54, 41, 76, 44]. While our sample consists of individuals with established schizophrenia, this finding raises the possibility that some patients might retain or revert to a NMDA hyperfunction typically associated with earlier stages of the illness. This could suggest a more dynamic and heterogeneous course of glutamatergic dysfunction in schizophrenia than previously thought.

#### 2. Compensatory Mechanism

The observed increase in NMDA self-connection could potentially represent a localized compensatory response to NMDA receptor hypofunction elsewhere in the brain. However, this interpretation is problematic. Previous studies have consistently shown NMDA hypofunction in primary sensory areas, including V1, in schizophrenia [14].

#### 3. Ketamine and Gamma Activity

The findings from Uhlhaas et al. (2018) [28] suggest that ketamine, an NMDA receptor antagonist, increased visual gamma activity, which contrasts with the decreased gamma activity typically observed in schizophrenia. This suggests that NMDA receptor dysfunction in schizophrenia may not simply involve hypofunction but could reflect a more complex or compensatory state, potentially explaining the increased NMDA-mediated excitation observed in our study.

#### 4. Schizophrenia sub-groups

The identification of potential subgroups within our cohort of individuals with schizophrenia suggested by our PCA could also explain the observed increase in NMDA-mediated excitation. This may reflect different neurophysiological profiles or trajectories of NMDA receptor function. One group may display the more classic NMDA hypofunction, while the other might exhibit an atypical pattern characterized by heightened excitation, possibly reflecting differences in disease progression, treatment response, or underlying genetic factors [78, 7]. It is also important to note that we did not test for copy number variations (CNVs), which are often associated with schizophrenia and could contribute to subgroup differences, as a high number of individuals with certain CNVs develop psychotic disorders [39, 46, 62].

The reduction in GABA-B-mediated inhibition we observed aligns with widespread reports of GABAergic dysfunction in schizophrenia [61, 30, 26]. This decrease in inhibitory control, combined with the unexpected increase in NMDA-mediated excitation, suggests a shift towards cortical hyper-excitability. This E/I imbalance is particularly significant in the superficial cortical layers, where the interplay between NMDA-mediated excitation and GABA-B-mediated inhibition is essential for various cognitive processes, including working memory and attention [8]. Their dysfunction could lead to prolonged disturbances in the fine-tuning of cortical excitability. When this inhibitory deficit is coupled with enhanced NMDA-mediated excitation, it potentially creates a scenario where neural circuits become overly responsive and less discriminating in their activation patterns.

### 4.3 Thalamic Projections and Negative Symptoms

We identified specific model parameters predictive of negative symptom severity through stepwise regression analysis (*F* = 7.74, *p* = 0.001, *r*^2^ = 0.51). Key predictors included AMPA-mediated connectivity from thalamic projection neurons to reticular neurons and GABA-B-mediated self-inhibition of superficial interneurons. These findings provide a potential link between circuit-level dysfunction and clinical manifestations of schizophrenia, particularly negative symptoms.

The involvement of thalamic projection neurons in predicting negative symptom severity is particularly interesting. Layer 6 cortical neurons projecting back to the thalamus play a critical role in modulating thalamic output and, subsequently, thalamocortical feedback [16]. Reduced efficacy in these projections could lead to diminished capacity to modulate and synchronize thalamo-cortical activity, potentially contributing to the cognitive and motivational deficits characteristic of negative symptoms [3]. The decreased modulation from layer 6 cortical neurons to the thalamus, coupled with altered inhibitory control from superficial interneurons, could lead to a dysfunctional thalamo-cortical loop. In this scenario, both the input to and output from the thalamus are not adequately regulated, potentially underlying the more pronounced negative symptoms seen in schizophrenia.

Our findings align with and extend the theory of thalamocortical dysfunction in schizophrenia. This hypothesis states that abnormal interactions between the thalamus and cortex contribute to the cognitive, perceptual, and emotional disturbances seen in schizophrenia [4, 5]. Specifically, it suggests that disrupted filtering and integration of sensory information by the thalamus leads to aberrant cortical processing and, consequently, to various symptoms of the disorder. This theory has been supported by a number of neuroimaging, electrophysiological, and post-mortem studies [82, 9, 18]. While previous research has highlighted the importance of thalamo-cortical interactions in schizophrenia, our work extends this by specifically implicating layer 6 cortical neurons projecting back to the thalamus, quantifying their role in negative symptom severity, and identifying the involvement of GABA-B-mediated self-inhibition of superficial interneurons.

### 4.4 Parameter restoration

Our parameter restoration analysis revealed a more complex picture of neural circuit alterations in schizophrenia than our initial group comparisons suggested. While our direct comparison of model parameters between schizophrenia and control groups identified only two significantly different parameters (increased NMDA-mediated recurrent excitation among superficial pyramidal cells and decreased GABA-B-mediated inhibition from superficial interneurons to superficial pyramidal cells), the restoration analysis involved adjustment of additional parameters.

To shift the spectral characteristics of the schizophrenia group to match those of healthy controls, we found that numerous parameters required adjustment, including:

- AMPA-mediated connections, particularly in thalamocortical circuits, which needed significant decreases.
- GABA-A mediated inhibition, requiring increases in superficial layers but decreases in deeper layers and thalamic projections.
- GABA-B mediated connections, needing diverse changes including increased self-inhibition of superficial pyramidal neurons.
- NMDA-mediated connections, generally requiring modest decreases across multiple pathways.

This discrepancy between the results of direct group comparisons and the restoration analysis highlights the non-linear nature of the relationship between model parameters and spectral outputs. It suggests that while individual parameters might not differ significantly between groups, their complex interactions and cumulative effects lead to the observed spectral differences in schizophrenia. This finding aligns with our model comparison results, which showed that the full model, incorporating all these receptor systems, best explains the observed spectral data in schizophrenia and healthy control.

## 5 Funding

This work was supported by CUBRIC and the Schools of Psychology and Medicine at Cardiff University andthe School of Psychology at University of Exeter. The data collection was funded by an MRC/EPSRC funded UK MEG Partnership Grant (MR/K005464/1) to KDS. ADS and LB are supported by a Wellcome grant (226709/Z/22/Z).

## 6 Acknowledgments

We thank Dr Laura Knight for data collection and Dr Daniel Hauke for proof reading and insightful comments on the manuscript.

## Notes

### Competing Interest Statement

The authors have declared no competing interest.

